# Stochastic microbiome assembly depends on context

**DOI:** 10.1101/2021.08.29.458111

**Authors:** Eric W. Jones, Jean M. Carlson, David A. Sivak, William B. Ludington

## Abstract

Observational studies reveal substantial variability in microbiome composition across individuals. While some of this variability can be explained by external factors like environmental, dietary, and genetic differences between individuals, in this paper we show that for the model organism *Drosophila melanogaster* the process of microbiome assembly is inherently stochastic and contributes a baseline level of microbiome variability even among organisms that are identically reared, housed, and fed. In germ-free flies fed known combinations of bacterial species, we find that some species colonize more frequently than others even when fed at the same high concentration. We develop a new ecological technique that infers the presence of interactions between bacterial species based on their colonization odds in different contexts, requiring only presence/absence data from two-species experiments. We use a progressive sequence of probabilistic models, in which the colonization of each bacterial species is treated as an independent stochastic process, to reproduce the empirical distributions of colonization outcomes across experiments. We find that incorporating context-dependent interactions substantially improves the performance of the models. Stochastic, context-dependent microbiome assembly underlies clinical therapies like fecal microbiota transplantation and probiotic administration, and should inform the design of synthetic fecal transplants and dosing regimes.

**Significance Statement:** Individuals are constantly exposed to microbial organisms that may or may not colonize their gut microbiome, and newborn individuals assemble their microbiomes through a number of these acquisition events. Since microbiome composition has been shown to influence host physiology, a mechanistic understanding of community assembly has potentially therapeutic applications. In this paper we study microbiome acquisition in a highly-controlled setting using germ-free fruit flies inoculated with specific bacterial species at known abundances. Our approach revealed that acquisition events are stochastic, and the colonization odds of different species in different contexts encode ecological information about interactions. These findings have consequences for microbiome-based therapies like fecal microbiota transplantation that attempt to modify a person’s gut microbiome by deliberately introducing foreign microbes.

## Introduction

The microbiome that organisms initially acquire tends to be stably maintained over time, with resulting physiological consequences for animal growth, development, and health [1, 2]. There is also substantial variation in microbiome composition between individuals: for example, the vast majority of bacterial species found in the human population are not present in a majority of humans [3, 4]. It is not yet known the extent to which this variability is driven by initial microbiome acquisition versus the subsequent ecological dynamics that occur once a species is stably colonized.

Colonization of the gut is stochastic, that is, exposure to a bacterial species does not guarantee colonization. Consequently, every time an organism encounters a bacteria, that bacteria might successfully colonize and begin to proliferate or it might not. These branching outcomes, generated continuously from every bacterial encounter of every organism, contribute to a baseline level of microbiome variability even among replicates in identical environments [5, 6]. The state of the microbiome can subsequently affect the odds that an invader species will successfully colonize—for example, with priority effects the establishment of one species may preclude or encourage subsequent colonization by another—and such feedbacks further complicate the microbiome assembly process [7, 8, 9].

Although diverse microbiome compositions are consistently observed in natural populations, there have been few attempts to systematically study the assembly of complex microbial communities under defined biological conditions and with known strains of bacteria [10, 11, 12]. Especially in vertebrates these “bottom-up” experiments are difficult to perform due to the sheer bacterial diversity of their gut microbiome, which can harbor on the order of 1,000 bacterial species (as in the typical human gut) [2]. Invertebrates, by contrast, often have simpler gut microbial communities: the microbiome of *D. melanogaster* contains on the order of 10 bacterial species with a core set of approximately five species [13, 14]. As in other animals the fly gut microbiome tends to be stable over time once initially colonized and has been linked to development, fecundity, and lifespan [15, 16]. Therefore, the fly gut microbiome exhibits variability across individuals, affects its host organism’s fitness, and constitutes a tractable experimental system that is representative of microbiome variability across organisms at large.

To probe how probabilistic colonization affects community assembly, we examined five core bacterial species of the *D. melanogaster* microbiome: *Lactobacillus plantarum* (LP), *Lactobacillus brevis* (LB), *Acetobacter pasteurianus* (AP), *Acetobacter tropicalis* (AT), and *Acetobacter orientalis* (AO). The genus *Acetobacter* consists of bacteria that metabolize various carbon sources including sugars, ethanol, and lactate and excrete acetic acid. Bacteria from the *Lactobacillus* genus metabolize amino acids and sugars and excrete lactic acid [17, 18, 19]. The yeast-based diet of *D. melanogaster* supplies bacteria in the gut microbiome with the nutrients needed to drive their ecological dynamics [20].

In this paper we characterize the microbiome assembly of *Drosophila melanogaster* and empirically show that these initial colonization events lead to microbiome variability in flies that are identically reared, housed, and fed. We took a combinatorial approach and inoculated each of the 31 combinations of the five core bacteria into separate groups of germ-free flies, with 48 fly replicates per combination (Fig. 1). The bacterial abundance of each species in each fly was assayed, and this abundance data was converted into presence/absence data to create a distribution of colonization outcomes for each bacterial combination. This data was previously published in Gould et al., PNAS 2018, but no analysis of stochastic microbiome assembly was performed at that time [16]. In this paper we first empirically characterize these colonization outcomes, then reproduce them with a sequence of increasingly complex mathematical models. We find that stochastic microbiome assembly generates variability in the fly gut microbiome, and that the colonization odds of each species are influenced by the context of the other species with which they are fed.

**Figure 1:**
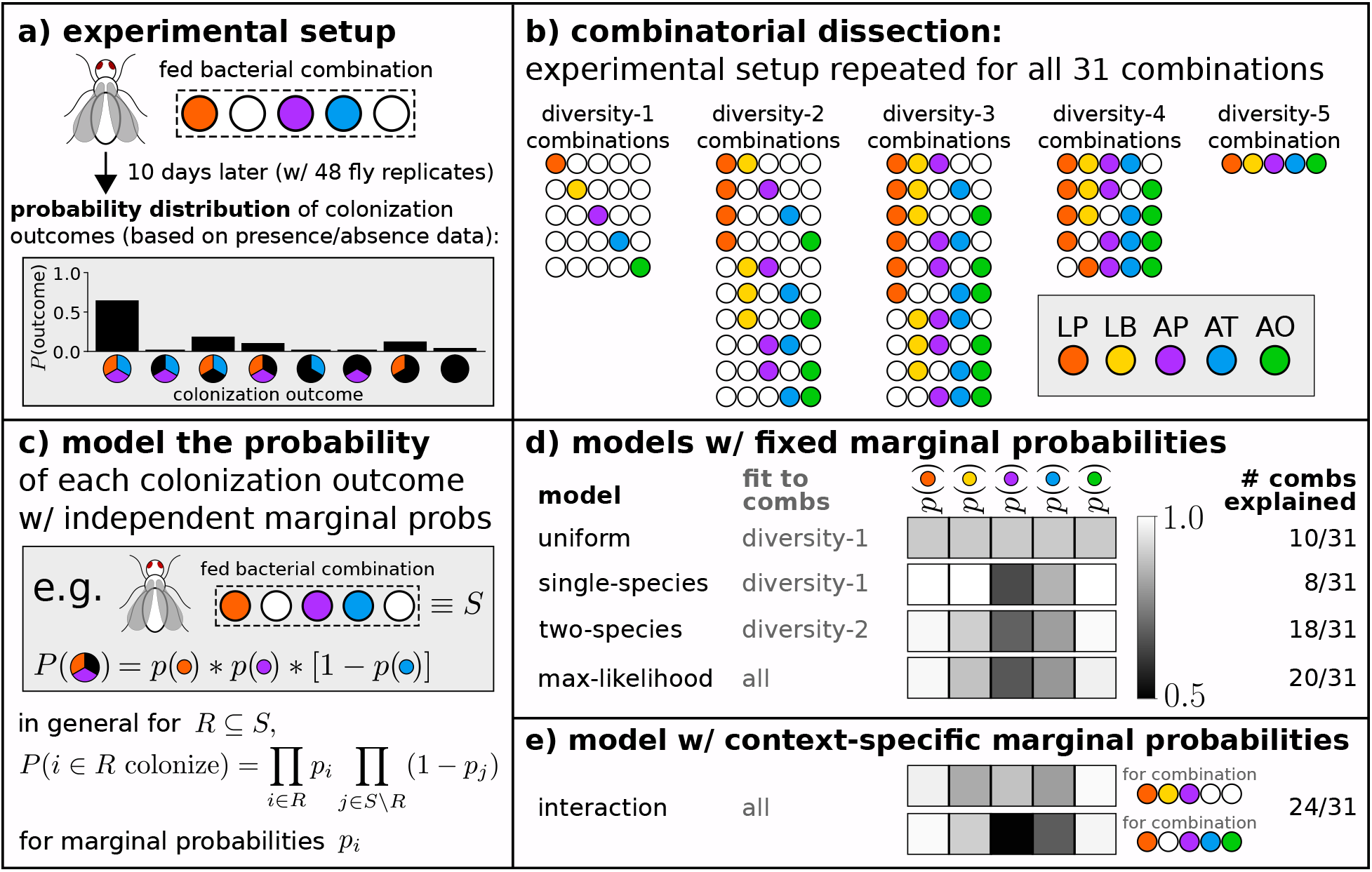
Experimental schematic and modeling framework. (a, b) Germ-free flies were associated with all 31 combinations of a core group of five bacteria, with 48 biological replicates per combination. Each fly was fed bacteria-laden food for 10 days, then crushed and plated to determine the bacterial abundance (measured as colony-forming units) of each species. Each species in a diversity-N combination can colonize or fail to colonize, yielding a probability distribution over 2^*N*^ colonization outcomes. Data previously published in Gould *et al*., *PNAS* 2018 [16]. (c) Empirical distributions of colonization outcomes (Fig. 2) are modeled assuming that the colonization of each species is independent, with species-specific colonization probabilities *P_i_* (also labeled *p*(*i*) in this schematic). (d) Models with fixed marginal colonization probabilities explain the colonization outcomes of up to 20 out of 31 bacterial combinations (multinomial test, *p* > 0.05); the two-species model performs nearly as well as the max-likelihood model but requires only a fraction of the data. (e) Models with context-specific marginal probabilities outperform models with fixed marginal probabilities.

## Results

### Standardized feeding leads to variable colonization outcomes

When a bacterial combination of *N* species (called a diversity-N combination) is fed to flies, 2^*N*^ colonization outcomes can result (corresponding to the presence/absence of each fed bacterial species). For each bacterial combination 48 fly replicates were fed bacteria-laden food in an identical manner, but some bacterial species failed to colonize some fly replicates which led to variable colonization outcomes (see Methods) [16]. Figure 2 plots the empirical frequency of each colonization outcome for each bacterial combination. The most common colonization outcome (for nearly every bacterial combination) is that all fed species colonize. Each additional fed species doubles the number of possible colonization outcomes; accordingly, at higher-diversity combinations more outcomes were observed.

**Figure 2:**
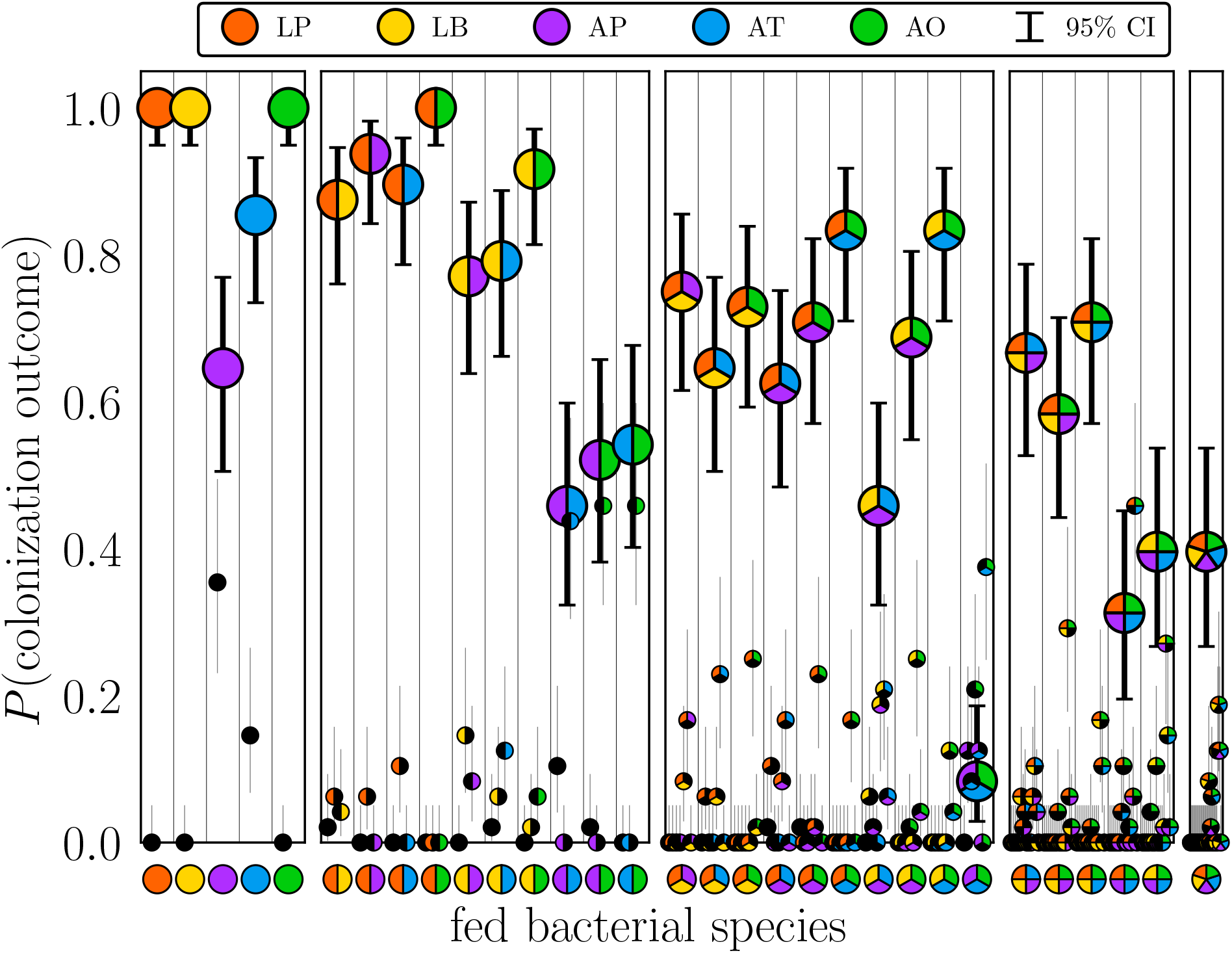
Distribution of colonization outcomes for each combination of fed species. Binary presence/absence data (detection limit of 100 bacterial CFUs) from 48 biological fly replicates per combination, represented as probabilities of colonization outcomes. Colonization outcomes are represented as pies with the number of slices equal to the combination diversity, colorful slices representing the presence of that bacterial species (as indicated in the legend), and black slices representing the absence of that species. Outcomes in which all fed species colonized are plotted as large pie markers. For each combination, the probabilities of all colonization outcomes sum to 1. Error bars indicate 95% confidence intervals of the mean.

Collapsing the colonization outcomes for experiments of each diversity yields the average number of species that successfully colonize as a function of the number of species fed, shown in Fig. 3a. Some species failed to colonize in some replicates for experiments of every diversity. Furthermore, as demonstrated in Fig. 3b the proportion of species that successfully colonize is relatively constant (ranging from 0.85-0.9) across combination diversities. Since variable colonization outcomes occurred even when flies were inoculated with bacteria at higher doses and in more uniform conditions than are typically found in nature, these findings suggest that stochastic colonization is a universal feature of microbiome community assembly.

**Figure 3:**
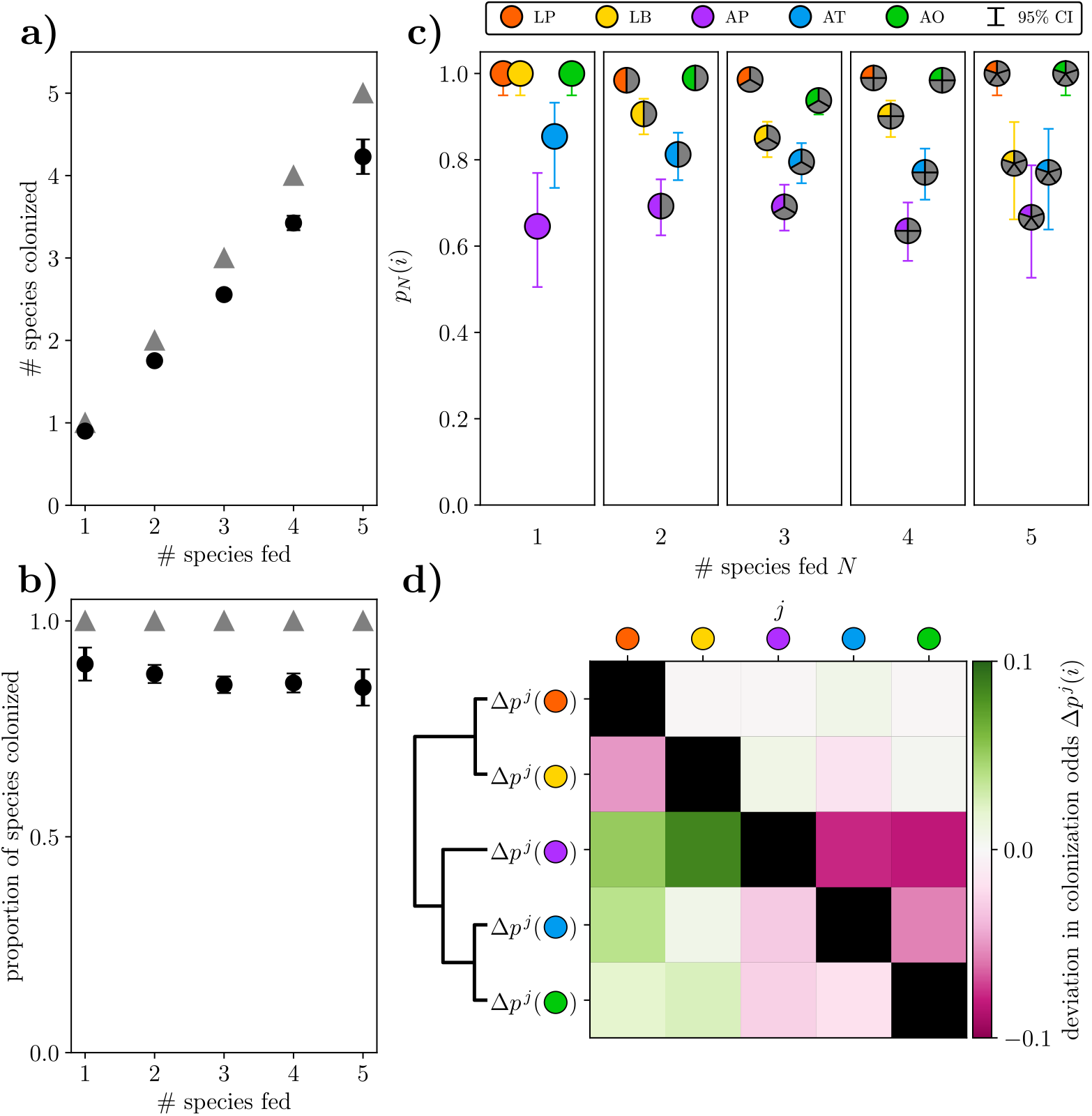
Empirical characterizations of colonization odds. (a,b) Bulk colonization properties quantify variability in colonization outcomes. Distributions of colonization outcomes (as in Fig. 2) are coarse-grained across species to yield (a) the average number of species that colonize and (b) the proportion of species that colonize for each combination diversity, plotted as black circles. Gray triangles indicate perfect colonization of every fed species. (c) Single-species colonization odds vary across combination diversities. For each bacterial species *i*, diversity-dependent colonization odds *p_N_*(*i*) were computed (across all experiments of diversity *N*) as the number of times a species successfully colonized divided by the number of times it was fed. Colorful slices represent the presence of that bacterial species (as indicated in the legend) and grey slices represent all outcomes for the other species. LB is less successful at colonizing in more diverse combinations (*p* = 0.02, Cochran-Armitage trend test). Error bars represent the 95% confidence interval of the mean. (d) Colonization odds depend on context. The deviation Δ*p^j^*(*i*) in the colonization odds of species *i* in the presence of species *j* is defined as the probability that species *i* colonized when fed with species *j* minus the probability that species *i* colonized (regardless of combination), and is plotted as a heatmap. *Acetobacter* species colonized more frequently in the presence of *Lactobacillus* species and less frequently in the presence of other *Acetobacter* species. Grouping the different rows of the heatmap by similarity yields the dendrogram at left, which accurately clusters bacterial species according to their genera.

### Colonization odds of bacterial species imply the existence of “strong” and “weak” colonizers

The colonization odds of each bacterial species—that is, the proportion of the time a bacterial species colonized when it was fed—differ in general, revealing a distinction between bacteria that are “strong colonizers” and others that are “weak colonizers.” Figure 3c shows the colonization odds *p_N_* (*i*) of species *i* in experiments of a given diversity *N*. These diversity-dependent colonization odds demonstrate that LP and AO are strong colonizers (colonizing more than 95% of flies they are fed to) while AT, and AP are relatively weak colonizers (colonizing less than 80% of flies they are fed to). The colonization odds of LB are 100% in single-species experiments but are significantly lower in higher-diversity combinations, possibly reflecting competitive exclusion of LB by other stronger colonizers when they are present.

### Species colonization odds depend on context

Bacterial species colonized with different odds depending on which other species were fed alongside. These context-dependent deviations in colonization odds Δ*p^j^*(*i*), defined as the colonization odds of species *i* in the presence of species *j* minus the colonization odds of species *i* regardless of combination, reflect interactions between bacterial species. Figure 3d shows a heatmap of these context-dependent deviations, indicating that *Acetobacter* species colonize more frequently in the presence of *Lactobacillus* species and less frequently in the presence of other *Acetobacter* species. The colonization odds of LP and LB are basically unaffected by the presence of other bacteria, while the colonization odds of AP, AT, and AO are sensitive to the presence of other bacteria. Clustering the rows of the heatmap yields the dendrogram at left, which correctly assigns bacterial species to their corresponding genera (the similarity metric comparing rows *i* and *j* only considers contributions from the three elements that are not *i* or *j*, see Methods). Notably, this taxonomic clustering is entirely based on distributions of colonization outcomes, meaning that in this case observational presence/absence data is sufficient to extract the functional similarity of species of the same genus.

### Reproducing empirical colonization outcomes with probabilistic models in which the colonization of each bacterial species is an independent stochastic process

Flies fed a diversity-*N* combination have 2^*N*^ colonization outcomes, corresponding to whether each species was able to colonize or not. We model these colonization outcomes by assuming that each species’ colonization is an independent process. More concretely, an *independent colonization model* posits that for a diversity-*N* combination *S*, the probability of a colonization outcome *R* ⊆ *S* is

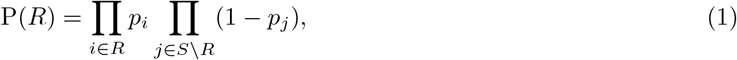

where *p_i_* is the marginal colonization probability of species *i*. Notationally we use “colonization odds” to refer to empirical quantities and “colonization probabilities” to refer to parameters of the independent models.

The choice of *p_i_* entirely determines the expected distribution of colonization outcomes of independent colonization models for a given bacterial combination. We consider two families of models: models with fixed colonization probabilities (*i.e*., models in which *p_i_* does not depend on the bacterial combination), and models with context-dependent colonization probabilities (*i.e*., models where *p_i_* can depend on the bacterial combination). We evaluate model performance with multinomial tests, by examining the likelihood of generating the observed data with a given model, and by computing the Bayesian Information Criterion (BIC) of each model (see Methods).

### Independent colonization models with fixed marginal colonization probabilities

We first consider four independent colonization models in which the colonization probabilities *p_i_* of each species are context-independent. Figure 1d shows a heatmap of *p_i_* in each of the four models, named the *uniform, single-species, two-species*, and *max-likelihood* models.

In the uniform model the colonization probabilities of each species are set to be identical, equal to the average colonization odds of single-species experiments. In the single-species model *p_i_* are set to the colonization odds of species *i* across the diversity-1 experiments; likewise, *p_i_* of the two-species model are set to the colonization odds of species *i* across diversity-2 experiments. Lastly, *p_i_* of the max-likelihood model are chosen to maximize the likelihood across experiments of all diversities, equally weighting the likelihood of generating the colonization outcomes of each combination. If the colonization probability of any species in any model is exactly 1 (as for LP, LB, and AO in the single-species model), we replace that probability by 0.9999 to avoid the possibility of colonization outcomes with zero likelihood, and in the Supplemental Information we show that the following results are not sensitive to this numerical choice.

It is pertinent for researchers to know whether independent colonization models with colonization probabilities *p_i_* fit to low-diversity (diversity-1 or diversity-2) experiments are similar to models fit to high-diversity (diversity-3 or higher) experiments, since if this is the case then a full combinatorial dissection is not needed to characterize the assembly of multi-species microbiomes. Figure 1d shows that colonization probabilities of the two-species model (which were fit to diversity-2 experiments) are very similar to the probabilities of the max-likelihood model (which were fit across every experiment). However, the probabilities of the single-species model differed: in diversity-1 experiments LB colonized 100% of the time, whereas in diversity-2 experiments it colonized only 91% of the time, and in higher-diversity experiments its colonization odds varied between 79% and 90%. These findings suggest that independent colonization models fit to two-species experiments implicitly account for bacterial interactions in a way that models fit to single-species experiments cannot.

### Independent colonization models with context-dependent marginal colonization probabilities

Next we consider a model in which the colonization probability *p_i_* of species *i* depends on the combination *S* in which it is fed (*i.e*., *p_i_* = *p_i_*(*S*)). The form of this *interaction model* is motivated by the context-dependent colonization odds in Fig. 3d. In the interaction model, when a *Lactobacillus* species *i* is fed with another *Lactobacillus* species, its colonization probability is adjusted so that 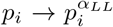. If a *Lactobacillus* species *i* is fed with an *Acetobacter* species, 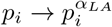, and if it is in the presence of both *Acetobacter* and *Lactobacillus* species, 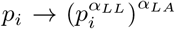. The same rules apply to *Acetobacter* species, but with parameters *α_AL_* and *α_AA_*. Exponentiation of colonization probabilities ensures that they remain bounded between 0 and 1. This particular model (with its relatively simple form) is representative of the improved performance of context-dependent colonization models broadly, although an infinite number of alternative context-dependent models might also demonstrate these properties.

The interaction parameters *α_LL_, α_LA_, α_AL_*, and *α_AA_* encode interactions between *Lactobacillus* and *Acetobacter* species. These parameter values were determined by maximizing the likelihood across all experiments, which yielded *α_LL_* = 2.4, *α_LA_* = 2.0, *α_AA_* = 2.6, and *α_AL_* = 0.5. Therefore, in the interaction model the colonization probabilities of *Lactobacillus* species are lower in the presence of any additional species. The colonization probabilities of *Acetobacter* species are lower in the presence of other *Acetobacter* species and are higher in the presence of *Lactobacillus* species. Weak colonizers are more strongly influenced by these interactions due to the form of the interaction model: for example, the colonization probability of AP decreases by 30 percentage points when it is in the presence of other *Acetobacter* species in the interaction model. The fit interaction parameters recapitulate the context-dependent deviation in colonization odds shown in Fig. 3d.

### Independent colonization models with context-dependent colonization probabilities reproduce empirical colonization outcomes better than models with context-independent colonization probabilities

The accuracy of independent colonization models is evaluated in two ways in Fig. 4. First, Fig. 4a shows when a model overestimates (green) or underestimates (pink) the probability that all fed species colonize. Second, Fig. 4b indicates when a model reproduces (white) or fails to reproduce (dark red) the observed colonization outcomes, as determined by the *p*-value of a multinomial test. The residuals of Fig. 4a capture model fits for the particular colonization outcome that all fed species colonize, while the *p*-values of Fig. 4b report deviations based on the entire observed and model-predicted distributions of colonization outcomes.

**Figure 4:**
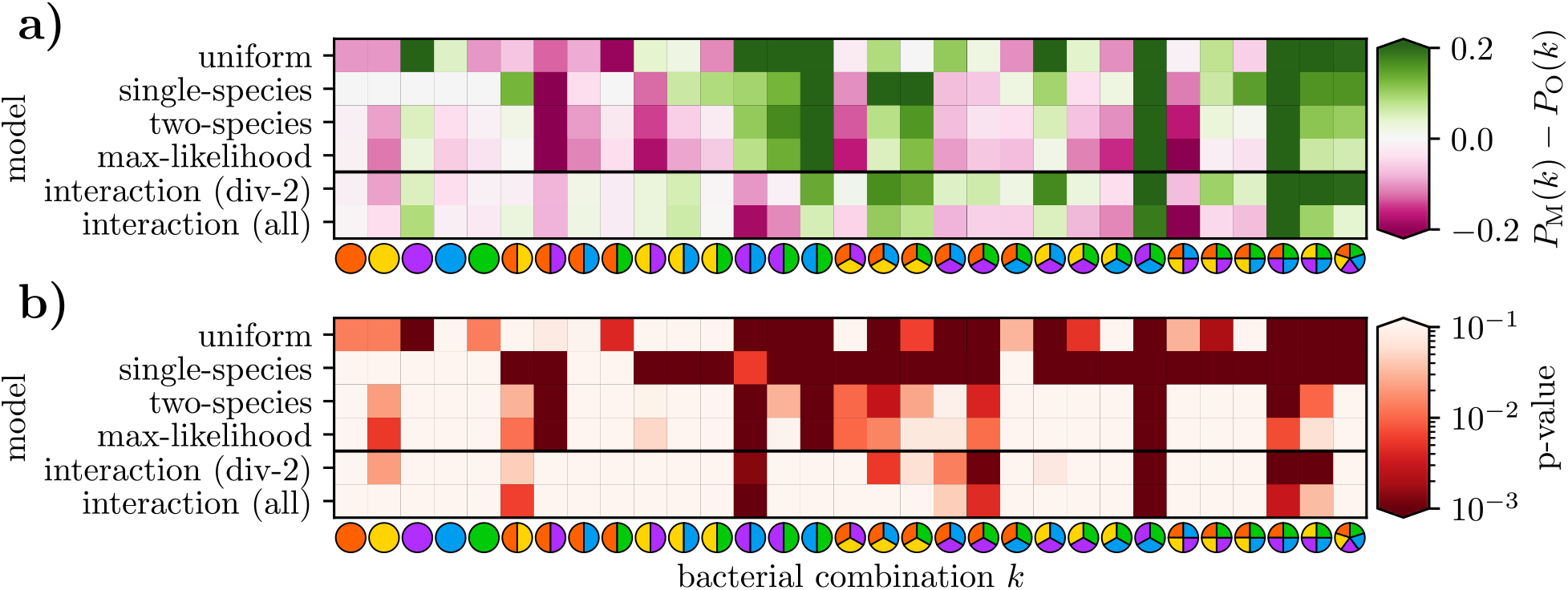
Evaluation of how independent colonization models reproduce empirical colonization outcomes. (a) Model residuals for the probability that all fed species colonize, where *P_M_*(*k*) and *P_O_*(*k*) are the model-predicted and empirically observed probabilities that all fed species of combination *k* successfully colonize, respectively. (b) The probability that a model reproduces the empirically observed distribution of colonization outcomes for each bacterial combination *k* (multinomial test; see Supplementary Information). In both panels, lighter colors indicate improved model fit, and the four models with fixed colonization probabilities are separated by a horizontal black line from the two models with context-dependent probabilities. The interaction model fit to all data is labeled “interaction (all)”, and the interaction model fit to diversity-2 experiments is labeled “interaction (div-2)”.

Figure 4a makes apparent that the context-dependent interaction models are better than models with fixed colonization probabilities at predicting the probability that all fed species colonize (*i.e.*, the bottom two models in each panel have lighter heatmap values than the top four rows). This trend is reinforced in Fig. 4b, where the context-dependent interaction models have higher *p*-values (*i.e.*, are better at reproducing the measured colonization outcomes) than models with fixed colonization probabilities.

Table 1 compiles additional performance measures for each model. The likelihood is defined as the probability of obtaining the observed data with a given model, in this case the product of the probabilities of 31 draws from multinomial distributions that each correspond to a particular combination of fed bacteria. The BIC is lowest for the interaction models. For two models *i* and *j* with equal priors, the quantity exp((BIC_*i*_ – BIC_*j*_)/2) roughly equals the odds that model *j* generated the observed data divided by the odds that model *i* generated the observed data [21]. As a rule of thumb, if the difference BIC_*j*_ – BIC_*i*_ > 10 then there is strong evidence to support model *i* rather than model *j*. Accordingly there is strong evidence to support models with context-dependent colonization probabilities over models with fixed colonization probabilities, and also to support models fit to all data over models fit to a subset of the data.

**Table 1:** Performance metrics for independent colonization models. The uniform model had one free parameter while the other models with fixed colonization probabilities had five free parameters. Both interaction models had nine free parameters, consisting of five colonization probabilities and four interaction parameters. Models that better combine empirical fit with fewer free parameters have smaller BIC scores.

| independent colonization model | # combinations reproduced ($p > 0.05$, multinomial test) | log-likelihood (across all experiments) | # free model parameters | Bayesian Information Criterion (BIC) |
| --- | --- | --- | --- | --- |
| uniform | 10/31 | -583 | 1 | 1169 |
| single-species | 8/31 | -1023 | 5 | 2063 |
| two-species | 18/31 | -331 | 5 | 680 |
| max-likelihood | 20/31 | -318 | 5 | 653 |
| interaction (div-2) | 22/31 | -298 | 9 | 626 |
| interaction (all) | 24/31 | -254 | 9 | 539 |

### Model failures indicate bacterial interactions

As is evident in Fig. 4b, the empirical colonization outcomes of most bacterial combinations are well-approximated by the independent colonization models. Nevertheless, these models fail to reproduce a few combinations (characterized by dark red vertical strips in Fig. 4b), and the models’ failure hints at an anomalous distribution of colonization outcomes. In two of these combinations, AP/AT and AP/AT/AO, the colonization odds of AP (45% and 33% respectively) are substantially lower than the average colonization odds of AP across all experiments (67%). Additionally, for AP/AT the colonization odds of AT (90%) are substantially higher than average (80%). These deviations indicate interactions between bacteria, in particular the exclusion of AP colonization by other *Acetobacter* species. The interaction model penalizes the colonization probabilities of *Acetobacter* species in the context of other *Acetobacter* species, yet it still failed to reproduce the distribution of colonization outcomes observed in AP/AT and AP/AT/AO. Partially, the interaction model is not compatible with the empirical observation that for the AP/AT combination the colonization odds of AT are higher in the presence of AP. More generally the rich structure of the combinatorial colonization outcomes lays bare the limitations of minimally context-dependent colonization models and motivates the future development of more intricate colonization models.

### Colonization odds inferred from diversity-2 experiments predict colonization outcomes of more diverse experiments

In experiments with multiple species the presence of bacterial interactions can affect colonization, and these effects can be accounted for with a more complex model. Taking these interactions into account—either by using the two-species model rather than the single-species model or by considering context-dependent colonization probabilities—improves the explanatory power of the associated independent colonization model. Of the independent models with fixed marginal colonization probabilities, the two-species model is nearly identical to the max-likelihood model, with the colonization probability of each species differing by ≤ 2 percentage points between the two models. Accordingly, these two models reproduce a similar number (18/31 and 20/31, respectively) of bacterial combinations and have similar log-likelihoods (−331 and −317). Notably, the single-species model performs much more poorly: when fed in isolation, LP, LB, and AO colonized perfectly, but their colonization odds worsened significantly once they were fed in combination with other species. Thus, considering bacterial context (and not using exclusively single-species experiments) significantly improves prediction. Supplemental Table 1 contains the log-likelihoods of each model for experiments of each diversity, which provides additional quantitative backing to this notion.

The context-dependent interaction model fit to diversity-2 experiments also outperformed (*i.e*., reproduced the colonization outcomes of more combinations, had a higher log-likelihood, and had a lower BIC score) all models with fixed colonization odds, including the max-likelihood model. The interaction model fit to diversity-2 experiments is sufficient to predict the majority (10/16) of high-diversity (diversity-3 and higher) colonization outcomes. For comparison, the interaction model fit to all experiments reproduces 11/16 of high-diversity colonization outcomes. Therefore interactions can be identified and accounted for based solely on two-species experiments.

## Discussion

### Stochastic colonization is a universal feature of microbiome assembly that produces microbiome variation across individuals

Microbiome assembly begins at birth, and the ecological dynamics that give rise to an individual’s microbiome can be separated into two sequential processes: probabilistic colonization by a bacterial inoculum, and subsequent internal dynamics (including replication, death, shedding, and secondary colonization by sloughed-off bacteria). By examining the stochastic colonization process in detail in the fruit fly, we demonstrate that a baseline level of microbiome variability exists among identically treated experimental replicates. In nature, bacterial inocula are typically less concentrated than they were in our experimental setup, so natural colonization events will therefore occur with lower odds than those that we observed, producing broader distributions of colonization outcomes than in our experiments.

### Stochastic microbiome assembly underlies bacteriotherapies like fecal microbiota transplantation (FMT) and probiotics

Treatments like fecal microbiota transplantation (FMT) and probiotics hinge upon successfully engrafting a “healthy” microbial community into a sick person’s microbiome [22]. While FMT is capable of producing long-term shifts in microbiome composition in some individuals, for others its effect can be muted or ineffectual [23]. Typical fecal transplants introduce hundreds of bacterial species into a sick person’s gut microbiome, but the imperfect colonization of these species might be partially responsible for the variable success of FMT treatments [24, 25, 26].

Currently, fecal transplant engraftment is typically quantified at a compositional level by examining how the donee’s microbiome changes over the course of a transplant (*e.g*., by measuring the species richness), and also by examining whether the donee’s microbiome becomes more similar (*e.g.*, as quantified by UniFrac) to the donor’s microbiome [27]. Quantifying the proportion of species from a fecal transplant that successfully engraft is complicated by the imperfect detection of low-abundance species, but some analyses of this type have nonetheless been performed. Le Roy, *et al*. implanted a fecal transplant containing 180 operational taxonomic units (OTUs; approximately equivalent to bacterial species) into germ-free mice, and after nine weeks 162 of these donor OTUs were found in the donee mice, so 90% of transplanted species successfully colonized [28]. Maldonaldo-Gomez *et al*. found the probiotic *Bifidobacterium longum* AH1206 in 30% of humans six months after ingestion [29]. These two observations imply that bacteriotherapies like FMT and probiotics are influenced by stochastic microbiome assembly.

Future predictive frameworks of FMT or probiotic efficacy should take into account the stochastic nature of community assembly. For instance, FMT “superdonors” (who provide fecal transplants that engraft especially frequently) might harbor an increased number of bacteria that are strong colonizers, as we observed in our experiments [22]. Additionally, given that some bacterial species will fail to successfully engraft during FMT, our framework points to the advantage of building synthetic fecal transplants with functional redundancy in mind so that each desired bacterial function can be performed by multiple types of bacteria present in the transplant. Previous work found that once a bacterial dose is high enough to saturate the dose-response curve (as was the case in our experiments), simply increasing the inoculation dose does not substantially affect that species’ colonization odds [5]. Inoculating with several functionally-similar bacterial species, on the other hand, takes advantage of the independent colonization odds of each species to improve the likelihood that at least one of the desired species will successfully colonize. Lastly, our framework can be straightforwardly extended to predict the probability that sequential FMT or probiotic administrations will be successful: for a bacterial species with per-treatment colonization odds *p_i_*, the probability it colonizes after N inoculations is 1 – (1 – *p_i_*)^*N*^. Thus, this framework could inform FMT and probiotic dosing regimens and lead to improved clinical outcomes.

### Presence/absence data reveals ecological interactions and taxonomic structure

We inferred interactions between pairs of bacteria by examining how the colonization odds of one species change when it is fed with the other species. Notably, these interactions were determined using only presence/absence data derived from bacterial abundance data. In natural environments with unknown exposures where organisms assemble their microbiomes from an environmental bath of bacteria, interactions between bacteria might be similarly identified using only this frequently collected presence/absence data. For example, in cohoused mice the context-dependent colonization odds of different species might reflect facilitative or competitive interactions between different species. Binary presence/absence data offers a particularly accessible lens for examining complex interactions in the microbiome.

### Two-species experiments predict colonization outcomes of higher-order experiments

Independent colonization models fit to diversity-2 experiments performed nearly as well as models fit to all experiments. Single-species experiments do not capture bacterial interactions, which limits their ability to explain the colonization behaviors of higher-order bacterial combinations. The colonization patterns of higher-order combinations are approximated reasonably well by independent colonization models in which the colonization odds of each species are determined with pairwise experiments, which is consistent with recent work studying multispecies synthetic soil microbiomes [10].

## Conclusion

Stochastic microbiome assembly is a ubiquitous process that occurs in all nascent microbiomes and has lasting consequences for microbiome composition. Our results detail variability in colonization outcomes in the fruit fly, even when flies are fed high doses of bacteria in identical experimental conditions. Our experiments sidestep the complication of historical contingency in community assembly (*i.e*., priority effects) by simultaneously feeding germ-free flies all types of bacterial species [7]. We find that context-dependent deviations in colonization odds reveal interactions between bacterial species: this method for inferring bacterial interactions is based on the ensemble of colonization outcomes across biological replicates, and differs from other traditional inference methods that attempt to infer interactions from compositional time-series data [30, 31]. Probabilistic models fit with low-diversity colonization data are capable of reproducing the colonization outcomes of higher-diversity combinations, which suggests that the acquisition of multi-species microbial communities can be coarse-grained in terms of the colonization odds of individual species. Stochastic microbiome assembly plays a crucial role in microbiome dynamics, and interrogating empirical colonization data with falsifiable mathematical models is a necessary step towards better understanding this process. Future efforts to deliberately drive an individual’s microbiome to a desired composition (the goal of personalized microbiome healthcare) will benefit from taking stochastic colonization into account when prescribing microbiome-based therapies.

## Materials and Methods

Additional details are provided in SI Appendix.

### Procedure for bacterial inoculation in germ-free flies

Data in this paper was published in Gould et al., PNAS 2018 [16]. Briefly, each of the 31 combinations of 5 core bacteria (LP, LB, AP, AT, and AO) were fed to 4 separate biological replicates of 12 germ-free flies (48 total flies per combination for 1488 total flies). 48 negative control flies were maintained germ-free. Vials of initially germ-free flies were inoculated with 5 * 10^6^ CFUs of each bacterial species. Flies consumed the bacteria and were transferred to vials with fresh bacteria-laden food every 3 days for 10 days. Individual flies were then surface-sterilized with 70% ethanol, crushed, and plated on agar plates to enumerate CFUs. Fig. S1 plots the distribution of CFUs for each species across all experiments. The abundance of stably colonized bacterial species was substantially higher than the limit of detection: median bacterial abundance of colonized flies was 152,000 CFUs; limit of detection was 100 CFUs. Therefore, while it is feasible that this presence/absence dichotomy excludes some flies that were minimally colonized, this occurrence appears to be relatively rare.

### Bacterial inoculation doses are at the “plateau” of the dose-response curve

In the fruit fly, the probability that bacterial strains colonize follows a dose-response curve that is an increasing function of the inoculum dose [5]. In the prior study by Obadia et al., single bacterial species were fed to flies in inoculum doses ranging from 10^1^ to 10^8^ CFUs: for example, a lab fly-gut isolate of *Lactobacillus plantarum* colonized 20% of flies when fed at a dose of ~5 CFUs, colonized 60% of flies when fed at a dose of ~300 CFUs, and colonized 70% of flies when fed at a dose of ~3,000,000 CFUs. This logistic shape—low colonization odds at low inoculum doses, with colonization odds plateauing (not necessarily at 100%) for high inoculum doses—was observed in all bacterial species (except for species that colonized 100% at all doses). Therefore, the colonization odds of individual species strongly depend on the inoculum dose, and are an important factor in determining colonization outcomes. To reduce the substantial variability in colonization outcomes of the fruit fly, we standardized our experimental procedure by fixing the inoculum size at 5 * 10^6^ CFUs for each bacterial species, and flies were permitted to feed continuously. This inoculum size saturates the dose-response curve so that higher doses do not lead to better colonization odds [5].

### Calculation of 95% confidence intervals of colonization odds

To compute the confidence intervals of Fig. 2, each colonization outcome was treated as a binomial variable (in which success was defined as that colonization outcome, and failure was defined as all other colonization outcomes), from which binomial confidence intervals were computed. Binomial confidence intervals do not take into account the covariance structure of multinomial proportions, and therefore are a conservative estimate of their confidence intervals. The binomial confidence intervals of Fig. 2 and Fig. 3abc were computed using the Jeffreys interval (derived from Bayesian statistics) as implemented in the statsmodels.stats.proportion.proportion_confint Python function.

### Multinomial tests, likelihood calculations, and Bayesian Information Criterion (BIC) scores

Each of the independent colonization models generates a predicted distribution of colonization outcomes that can be compared to the empirical colonization outcomes using multinomial tests, likelihood calculations, and BIC scores. For each bacterial combination, multinomial tests use the multinomial distribution of colonization outcomes generated by an independent model as a null model and calculate how likely it is for this null model to generate the observed multinomial distribution of colonization outcomes or a less likely distribution (see Fig. S1 for a schematic explanation). Exact multinomial tests (used for diversity-1 and diversity-2 combinations) or Monte Carlo multinomial tests (used for diversity-3 and higher combinations) were computed with the XNomial package in R. Likelihood computations measure the probability that this same null model would yield exactly the observed distributions of colonization outcomes, *i.e*., the probability of “drawing” the empirical distribution of outcomes with that multinomial null model. Lastly, the Bayesian information criterion (BIC) is a metric that quantifies the trade-off between increased model accuracy and increased complexity by rewarding models with a high likelihood and penalizing models with more free parameters: BIC = *k* log *n* – 2 log *L*, where *k* is the number of free parameters of the model, *n* is the sample size, and *L* is the likelihood of the observed data given the model.

## Software availability

The software and raw data used to perform analyses and generate figures in this study is available online on GitHub (https://github.com/erijones/stochastic_microbiome_assembly).

## Acknowledgements

E.W.J. was supported by Banting and Pacific Institute for the Mathematical Sciences Postdoctoral Fellowships. The David and Lucile Packard Foundation and the Institute for Collaborative Biotechnologices suppported J.M.C. through Grant W911NF-09-0001 from the US Army Research Office. D.A.S. was supported by a Natural Sciences and Engineering Research Council of Canada Discovery Grant and by the Canada Research Chairs program. W.B.L. was supported by National Institutes of Health grant DP5OD017851, National Science Foundation Integrative Organismal Systems award 2032985, and the Carnegie Institution for Science Endowment. E.W.J., D.A.S., and W.B.L. were supported by a Carnegie Institution of Canada grant.

